# How the terminal glucoside of the N-glycan donor affects the catalytic efficiency of the eukaryotic oligosaccharyltransferase

**DOI:** 10.64898/2026.07.16.738906

**Authors:** Beatrice Tropea, Elisa Fadda

**Affiliations:** Hamilton Institute, Maynooth University, Maynooth, Ireland; School of Biological Sciences, University of Southampton, Southampton, United Kingdom

## Abstract

The eukaryotic oligosaccharyltransferase (OST) is the enzyme responsible for initiating N-glycosylation of secreted proteins by transferring a pre-assembled lipid-linked oligosaccharide (LLO) donor to target asparagine residues most often found within N-x-S/T consensus sequences, or sequons. OST preferentially selects LLO donors with a distinctive glucoside Glc-α(1-2)-Glc-α(1-3)-Glc-α(1-3)-capping the A-branch. After the N-glycosylation reaction, this motif is cleaved in a stepwise manner from the immature N-glycan structure before the folded glycoprotein exits the endoplasmic reticulum quality control (ERQC) cycle. While the -Glc-α(1-3)-Glc-α(1-3)-disaccharide is an important flag regulating binding to the calreticulin/calnexin chaperones, the terminal Glc-α(1-2)-is removed immediately after OST catalysis, suggesting that its biological function may be directly linked to the OST catalytic efficiency. To understand how and why this capping motif affects the OST N-glycosylation efficiency, we rebuilt 3D models of the yeast OST in complex with an acceptor peptide and LLO donors substrates with and without terminal Glc-α(1-2)-, and analysed their stability and dynamics with all-atom molecular dynamics (MD) simulations through both conventional, and Gaussian-accelerated (GaMD) sampling schemes. Our results indicate that the terminal Glc-α(1-2)-is essential to anchor the full-length LLO donor to the OST through a complex network of intermolecular contacts extending from the catalytic site to distal subdomains. We show how this contact network is crucial to preserve the LLO catalytically productive alignment of its reducing end. We also show that the removal of the terminal Glc-α(1-2)-leads to an increased flexibility of the LLO, which displaces the reducing end and redistributes the conformational ensemble towards misaligned states, which are less catalytically productive. These results provide a mechanistic basis linking the catalytic efficiency of the eukaryotic OST to the distinctive glucosylated structure of the LLO donor.

## Introduction

N-glycosylation is one of the most common post-translational modification (PTM) of proteins, and a key regulatory mechanism in eukaryotic cells mediating folding, stability, trafficking and cell signalling (Aebi 2013; Cherepanova, Shrimal and Gilmore 2016; Varki *et al*. 2022). Mammalian nascent proteins entering the secretory pathway are co- or post-translationally N-glycosylated in the ER lumen by two oligosaccharyltransferase complexes, namely OST-A and OST-B(Kelleher *et al*. 2003; Mueller *et al*. 2015; Shrimal, Cherepanova and Gilmore 2015; Ramírez, Kowal and Locher 2019; Ramírez and Locher 2023), respectively. Both complexes have largely similar architectures and domains sequences(Ramírez, Kowal and Locher 2019), with OST-A operating at the translocon(Shrimal, Cherepanova and Gilmore 2017; Gemmer *et al*. 2023) and responsible for the majority of protein N-glycosylation reactions, with OST-B selecting some of the sites skipped by OST-A(Ruiz-Canada, Kelleher and Gilmore 2009). In this work we focus on the major *S. cerevisiae* Ost3-OST (yeast OST)(Schulz *et al*. 2009) corresponding to the OST-B for which a cryo-EM structure of the ternary complex is available(Ramírez *et al*. 2022), where the yeast OST is bound to a non-reactive peptide acceptor and to a full LLO substrate donor.

The OST catalytic function involves the transfer of a fully assembled Glc_3_Man_9_GlcNAc_2_ oligosaccharide linked to a dolichol pyrophosphate, known as lipid-linked oligosaccharide (LLO), to specific asparagine residues most commonly found within a consensus sequence N-x-S/T (where x can be any amino acid but proline), known as sequon (Marshall 1972; Zielinska *et al*. 2010; Aebi 2013; Cherepanova, Shrimal and Gilmore 2016). OST is a large multimeric membrane protein complex (**Fig. 1.a**), composed of eight subunits arranged into three functional groups(Bai *et al*. 2018; Wild *et al*. 2018; Ramírez *et al*. 2022), subcomplex I (Ost1 and Ost5), subcomplex II, which contains the catalytic subunit Stt3 together with Ost4 and either Ost3 or Ost6, and subcomplex III (Ost2, Wbp1, and Swp1). While the catalytic Stt3 is conserved across all domains of life, in eukaryotes auxiliary subunits evolved to enhance substrate recognition and catalytic efficiency (Schwarz, Knauer and Lehle 2005; Schulz *et al*. 2009; Bai *et al*. 2018; Wild *et al*. 2018; Eyring *et al*. 2021; Neuhaus *et al*. 2021).

**Figure 1.**
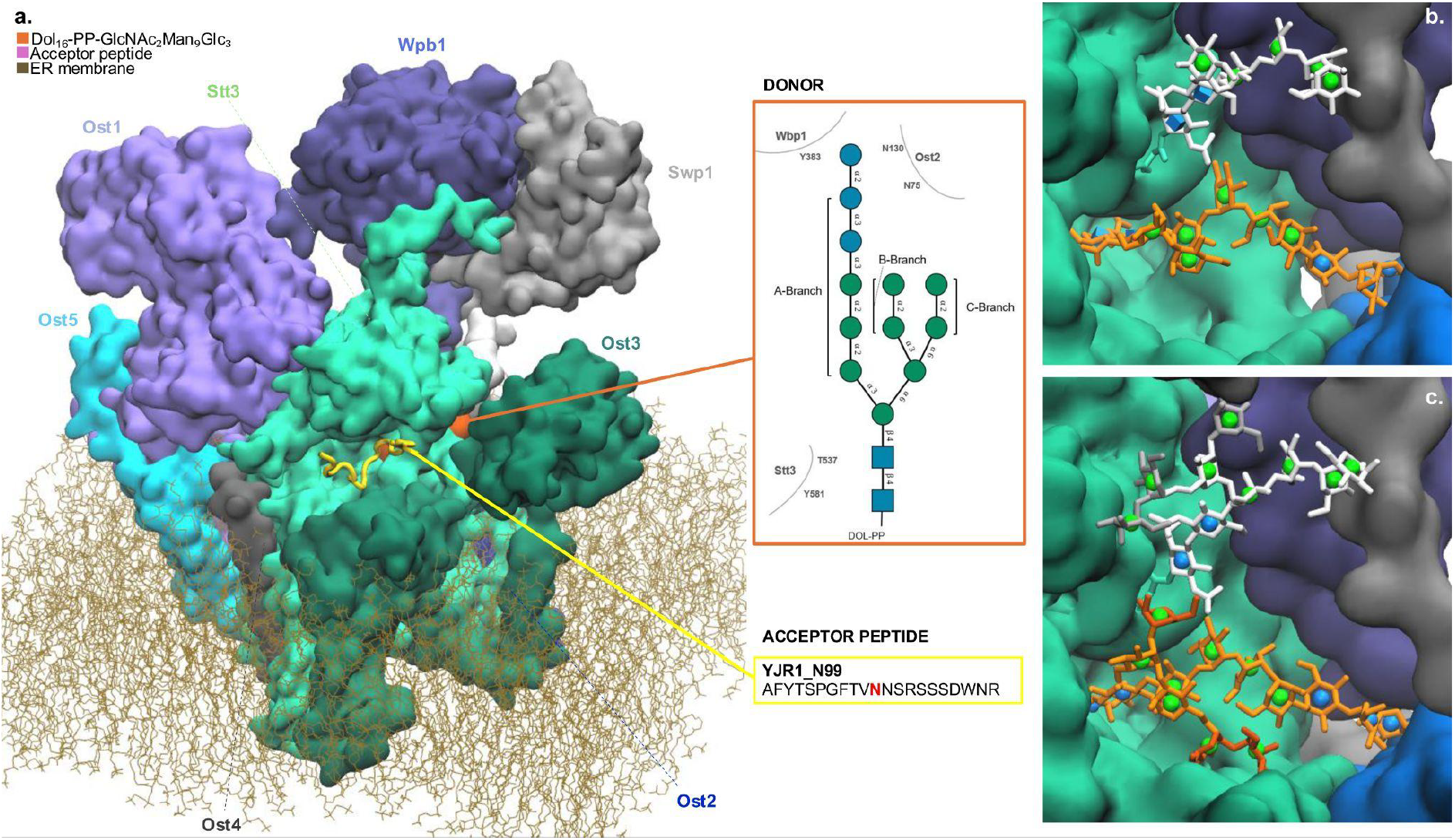
Structural overview of the yeast OST complex and donor-binding site. **(a)** 3D structure of the yeast OST complex embedded in the ER membrane (brown sticks), showing the 8 subunits as surface color coded according to the labels. The lipid-linked oligosaccharide (LLO) donor is shown in orange and the backbone of the acceptor peptide is shown with a cartoon representation in yellow. The central panel shows the 2D SNFG representation of the LLO donor, with labels specifying glycosidic linkages and the nomenclature used for the branches. Below we show the sequence of an acceptor peptide, i.e. YJR1_N99, used as substrate in all MD simulations in this work, where the target asparagine was previously characterised as efficiently glycosylated(Khaleque *et al*. 2025). **(b)** Close-up view of the donor-binding pocket from the cryo-EM structure (PDB ID: 8AGC), showing the resolved LLO donor represented in sticks (orange) and SNFG symbols and the Stt3 N539 glycan (white). **(c)** Close-up view of the donor-binding pocket from the 3D model used in the MD simulations with the reconstructed N-glycans. The monosaccharides resolved in the cryo-EM structure are shown in white and orange as in panel b, whereas the monosaccharides added in the reconstruction of the full LLO donor and N539 glycan are highlighted in dark orange and grey, respectively. Rendering with Visual Molecular Dynamics (Humphrey, Dalke and Schulten 1996) (VMD) (https://www.ks.uiuc.edu/Research/vmd/).

A critical aspect of the eukaryotic N-glycosylation process is the requirement for a fully assembled LLO donor structure Glc_3_Man_9_GlcNAc_2_ containing three terminal glucoses (Burda and Aebi 1998; Burda *et al*. 1999; Karaoglu, Kelleher and Gilmore 2001; Kelleher *et al*. 2003; Alexander *et al*. 2026) (**Fig. 1.a**). The removal of the capping glucoside proceeds rapidly after OST catalysis through the activity of α-glucosidase I (GI) and α-glucosidase II (GII), so that none of the glucose units are ever detected in secreted proteins, except in rare contexts where the N-glycosylation site is structurally inaccessible (Crispin *et al*. 2004; Zhao *et al*. 2022). While GI removes the terminal Glc-α(1-2)-immediately after N-glycan transfer(Burda P 1998; Deprez, Gautschi and Helenius 2005), GII removes the α(1-3)-linked glucoses in a concerted manner within the calreticulin/calnexin (Cnx/Crt) cycle as part of the ER quality control (ERQC) system (Molinari and Helenius 2000; Deprez, Gautschi and Helenius 2005; Cherepanova, Shrimal and Gilmore 2016; Guay *et al*. 2025). Earlier work showed that the absence of the terminal glucoses on the LLO substantially reduces turnover rates (Burda *et al*. 1999; Haeuptle and Hennet 2009; Hennet 2012; Zacchi and Schulz 2016; Poljak *et al*. 2018), indicating the incorporation of this moiety contributes directly to the OST catalytic efficiency.

Recent structural insight from cryo-EM on the *S. cerevisiae* OST ternary complex(Ramírez *et al*. 2022) reveals a key pocket region at the Wbp1–Ost2 interface where the glucoside moiety resides. This pocket also bears a highly conserved N-glycan at N539 of Stt3 (**Fig. 1.c** and **1.d**)(Wild *et al*. 2018; Shrimal and Gilmore 2019). This detailed structural insight provides us with the ideal framework to investigate at the atomistic level how and why the LLO terminal glucoside is determinant for OST catalytic efficiency, and how the alteration of this motif reshapes the donor’s conformational landscape. Indeed, the misalignment of the donor’s reducing end relative to the target asparagine side chain on the acceptor peptide and to the catalytic residues would negatively affect the N-glycosylation efficiency (Lairson *et al*. 2008; Lizak *et al*. 2013; Ramírez and Locher 2023; Khaleque *et al*. 2025). In this study we use molecular dynamics (MD) simulations, using a combination of deterministic (conventional) and Gaussian accelerated (GaMD) sampling techniques, to investigate how the terminal Glc-α(1-2)-of the glucoside moiety stabilises the donor in a catalytically productive conformation in the yeast OST ternary complex. Through a comparison of the results obtained from the MD simulations of the OST in complex with the full-length donor, ie. Glc3Man9GlcNAc2-PP-Dol, with those obtained with a truncated intermediate lacking the terminal glucose, ie. Glc2Man9GlcNAc2-PP-Dol, we show that the capping Glc-α(1-2)-unit on the A-branch of the LLO is critical to stabilise its structure in a catalytically productive conformation. We show that the removal of the capping α(1-2)-Glc leads to a higher flexibility of the A-branch and to a displacement of the GlcNAc at the reducing end from the optimal productive state.

Previous studies addressed the link between donor structure and OST catalysis through genetic knockout of Alg6, which transfers the first Glc unit on the A-branch of the Man9GlcNAc2- (Reiss *et al*. 1996; Zacchi and Schulz 2016; Eyring *et al*. 2021; Ramírez *et al*. 2022; Khaleque *et al*. 2025). Mutations to the *ALG6* gene have been also linked to congenital disorders of glycosylation (CDG)(Westphal *et al*. 2000; Haeuptle and Hennet 2009), while mutations to *ALG10*, which encodes the enzyme that transfers the capping Glc-α(1-2)-, were reported only recently as linked to epilepsy, brain atrophy, and sleep abnormalities in humans(Gill *et al*. 2024). In this work we focus on the effect of lack of the terminal Glc-α(1-2)-as it was shown to be mechanistically linked directly to the OST catalytic activity(Burda and Aebi 1998; Zacchi and Schulz 2016). Earlier work in plants also shows that the lack of terminal Glc-α(1-2)-reduces the overall degree of N-glycosylation and negatively impacts normal leaf growth(Farid *et al*. 2011). Our atomistic and dynamic perspective supports these conclusions and demonstrates how the terminal α(1-2)-Glc-in the LLO structure is required to stabilise the donor’s catalytically productive conformation within the context of the yeast OST architecture(Kelleher *et al*. 2003), explaining the preference of the eukaryotic OST for a fully glucosylated LLO and its role in optimizing N-glycosylation efficiency.

## Results and Discussion

### The LLO structure complements the architecture of the OST donor binding pocket

To investigate how the structure of the LLO modulates OST catalysis, we rebuilt two 3D models of the yeast OST ternary complex, one with the LLO structure complete, and the other with the LLO missing the terminal α(1-2)-Glc-on the A-branch. Both 3D models were built from cryo-EM data (PDB 8AGC) (Ramírez *et al*. 2022), where we included explicitly only the OST subunits directly involved in donor binding, namely Stt3 (aa 6–682), Ost3 (aa 211–345), Ost2 (aa 22–130), Swp1 (aa 163–283), and Wbp1 (aa 263–418) (**Fig.1.a**). In both models we replaced the non-reactive acceptor in the cryo-EM structure with a 20 aa acceptor peptide from a yeast cell wall protein, namely YJR1_N99 (**Fig.1.a** and **Fig. 2.a** and **2.b**), that we demonstrated in earlier work to be consistently glycosylated(Khaleque *et al*. 2025). The structures of the conserved N539 glycan, modelled here as a Man8GlNAc2-(Li *et al*. 2005), and of the LLO donor were largely resolved by cryo-EM (**Fig.1.b**) and we retained this information in our model, only completing the 3D structures with the missing monosaccharides (**Fig.1.c**). We analysed the structure stability and dynamics of these two OST ternary complexes by MD simulations using initially a deterministic (conventional) sampling scheme for 2 μs of production for each system. Further details on the simulation set-up and protocols are included in the Computational Methods section.

**Figure 2.**
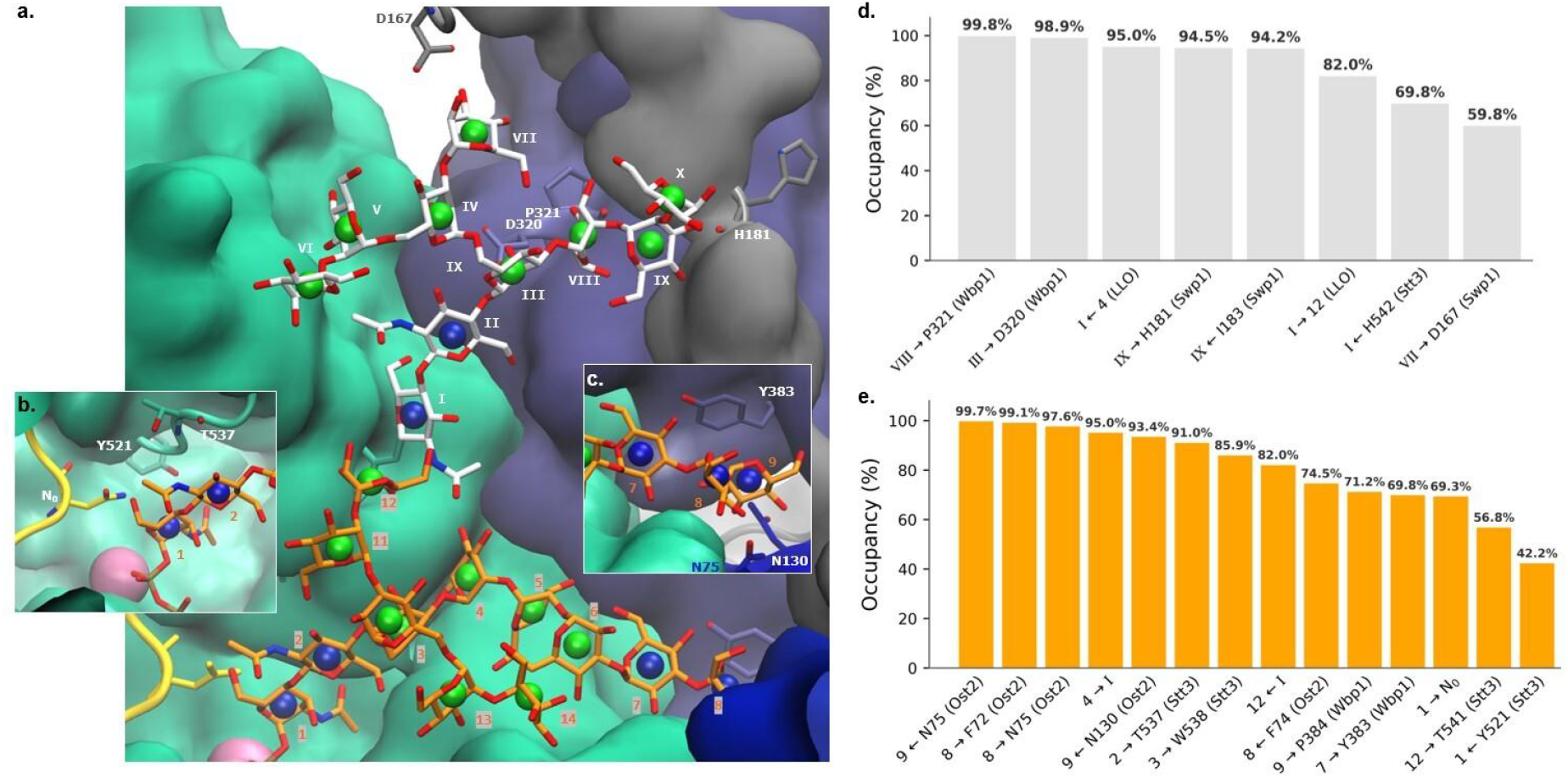
Interactions of the donor LLO and the N539-linked glycan with the OST complex. **(a)** Structure of the OST complex from the MD simulation showing the donor Glc_3_Man_9_GlcNAc_2_-PP-Dol (orange) and the N539 glycan modelled as a Man_8_GlcNAc_2_ (GlyTouCan ID G48369JO in white) according to ref.(Li *et al*. 2005), together with their corresponding SNFG representations. The protein surface is coloured by subunit, and OST residues involved in glycan binding are shown as sticks. **(b)** Close-up of the catalytic region showing interactions between the reducing end of the donor glycan, the acceptor sequon centred on N0, and residues T537 and Y521 involved in donor binding. **(c)** Close-up of the terminal Glc_3_ branch showing interactions with Asn75 and Asn130 of Ost2 and Tyr383 of Wbp1. **(d)** Hydrogen bonds analysis of the conserved N539-linked glycan throughout the MD simulation. **(e)** Hydrogen bonds analysis of the Glc_3_Man_9_GlcNAc_2_-PP-Dol donor throughout the simulation. Only interactions with an occupancy ≥40% are shown. Rendering with Visual Molecular Dynamics (Humphrey, Dalke and Schulten 1996) (VMD) (https://www.ks.uiuc.edu/Research/vmd/).

During the MD trajectory of the OST in complex with the intact LLO, the glucoside moiety remains stably anchored to residues in a pocket at the Wbp1–Ost2 interface (**Fig. 2.c** and **2.e**). The terminal Glc-α(1–2)-, indicated as ‘9’, forms a CH–π interaction with the Y383 (Wbp1) sidechain, while the OH of C3 engages in a hydrogen bond with the sidechain of N130 (Ost2). The sidechain of N75 (Ost2) coordinates the adjacent -Glc-α(1–3)-unit (‘8’) (**Fig. 2.c**). The conserved N539 glycan interacts with residues in both Wbp1 (purple), and Swp1 (grey) subunits **(Fig. 2.a)**, framing it in the donor-binding site. More specifically, the OH of C3 of the terminal Man_(X)_-α(1–2)-on the A-branch interacts with the backbone carbonyl of H181 in Swp1, while the -Man_(VIII)_-α(1–3)-engages the backbone carbonyl of P321 in Wbp1 **(Fig. 2.d)**. The central -Man_(III)_-β(1–4)-of the core pentasaccharide also forms a stable contact with the sidechain of D320 in Wbp1. The GlcNAc at the reducing end of the N539 glycan also contributes to restraining the LLO donor by forming an hydrogen bond with the -Man_4_-α(1–3)-of its A-branch. During the whole trajectory, the LLO -GlcNAc-β(1-4)-GlcNAc-core remains engaged to Stt3 through interaction with residues T537 and Y521 **(Fig. 2.b)**, as reported in the cryo-EM structure(Ramírez *et al*. 2022).

Taken together, these MD simulations results show that the structure of the LLO complements the architecture of the donor binding pocket in the OST, where specific interactions with the protein residues and with the conserved N539 glycan stabilize the catalytically productive orientation of the LLO in the catalytic site through the anchoring of the terminal Glc-α(1–2)-on the distal Wbp1–Ost2 pocket. This interaction network contributes to restrain the LLO inherent flexibility (**Fig.3**), reducing its conformational heterogeneity. These results are also in line with the structural analysis by cryo-EM where it was possible to clearly resolve most of the LLO structure(Ramírez *et al*. 2022).

### Removal of the terminal Glc-α(1–2)-triggers a change of coordination of the LLO in its binding pocket

When the LLO donor structure lacks the terminal Glc-α(1–2)-, the pocket at the Wbp1–Ost2 interface remains unoccupied, resulting in the loss of all the interactions that contribute to stabilise the A-branch (**Fig. 3.a**). The MD simulation results show that the removal of the terminal Glc-α(1–2)-determines a shift of the shorter the A-branch towards this distal pocket to re-establish the lost contacts. This event is coupled to a displacement of the GlcNAc at the reducing end relative to its position in the full-length system (**Fig. 3.b**), which disrupts the optimal catalytic orientation; more to this below. The deterministic MD simulation shows that the loss of distal anchoring is compensated by alternative contacts that stabilize the truncated donor into a different binding mode. These include interactions with Stt3 residues N536 and Q253, and Wbp1 residues H77 and N380 (**Fig. 3.c**). These interactions contribute to stabilising the truncated LLO donor within the binding pocket (**Fig. 3.d** and **3.e**), but the displacement at the catalytic site likely corresponds to a less productive binding mode.

**Figure 3.**
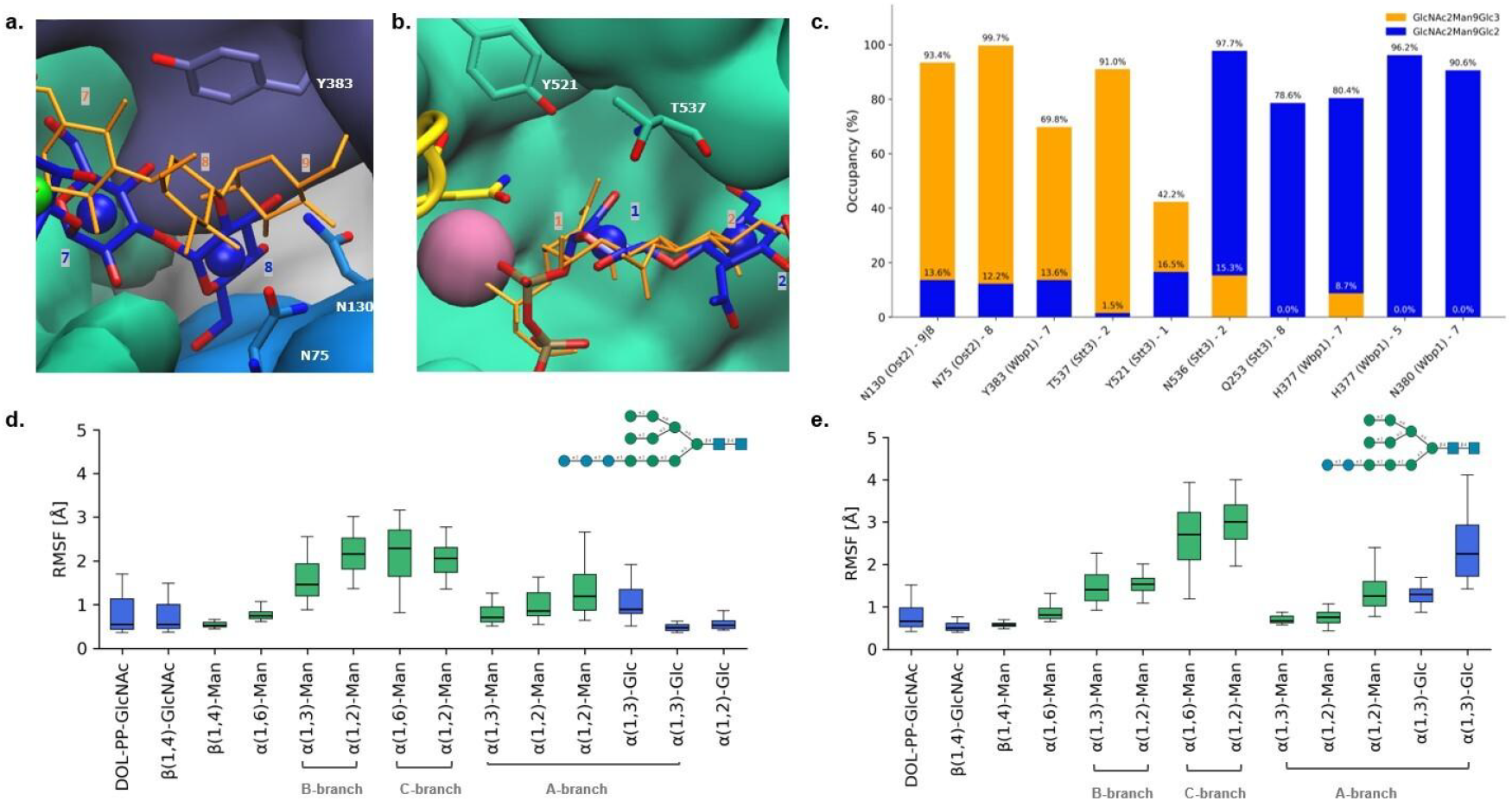
Comparison of donor binding in full-length and truncated LLOs. **(a)** Comparison of pocket occupancy between full-length (orange sticks) and truncated donors (blue sticks). In the full-length donor, the terminal glucose moiety occupies the pocket and stabilises local interactions, whereas in the truncated donor, the pocket remains unfilled, leading to rearrangements around residues Y583, N130, and N75. **(b)** Structural alignment of the full-length donor (orange) and the truncated donor (blue), showing that removal of the terminal glucose displaces the catalytic GlcNAc from a transfer-competent configuration. Residues Y521 and T537 are shown for reference. Rendering with Visual Molecular Dynamics (Humphrey, Dalke and Schulten 1996) (VMD) (ks.uiuc.edu/Research/vmd). **(c)** Hydrogen bond occupancy (%) for key donor–protein interactions in the full-length (orange) and truncated (blue) donors. The full-length donor maintains high-occupancy contacts that define the catalytic interface, whereas truncation disrupts these interactions and promotes alternative contacts. **(d**,**e)** Root-mean-square fluctuations (RMSF) values of individual donor monosaccharides for Glc_3_Man_9_GlcNAc_2_-PP-Dol (d) and Glc_2_Man_9_GlcNAc_2_-PP-Dol (e). Monosaccharides are grouped according to the A-, B- and C-branches of the donor glycan as indicated in **Fig. 1**.

### The terminal Glc-α(1–2)-is required to preserve a transfer-competent geometry

To test the conformational stability of the truncated LLO, we supplemented the conventional MD discussed above with enhanced sampling using a Gaussian Accelerated MD (GaMD) scheme. GaMD augments conformational exploration and allows us to access potential structural rearrangements that may be kinetically inaccessible under conventional sampling conditions (Miao, Feher and McCammon 2015; Wang *et al*. 2021). The GaMD results show that the truncated donor can indeed access four different conformers within the binding pocket (**Fig. 4.a-c**), consistent with the weakly restrained alternative binding interactions we identified through conventional MD sampling. Clustering analysis shows that the dynamics of the truncated LLO donor is largely driven by the A-branch not anchored through in the pocket at the Wbp1–Ost2 interface. This flexibility also enhances the dynamics at the reducing end, which affects the relative position and orientation of the anomeric carbon C1 relative to the catalytic residues (**Fig. 4.a**). Our analysis also shows that the interconversion between conformers across the trajectory is quite uniform (**Fig. 4.c**), with all four clusters significantly populated during sampling, indicating that the truncation of the LLO at the capping Glc-α(1–2)-is likely detrimental towards the OST catalytic efficiency.

**Figure 4.**
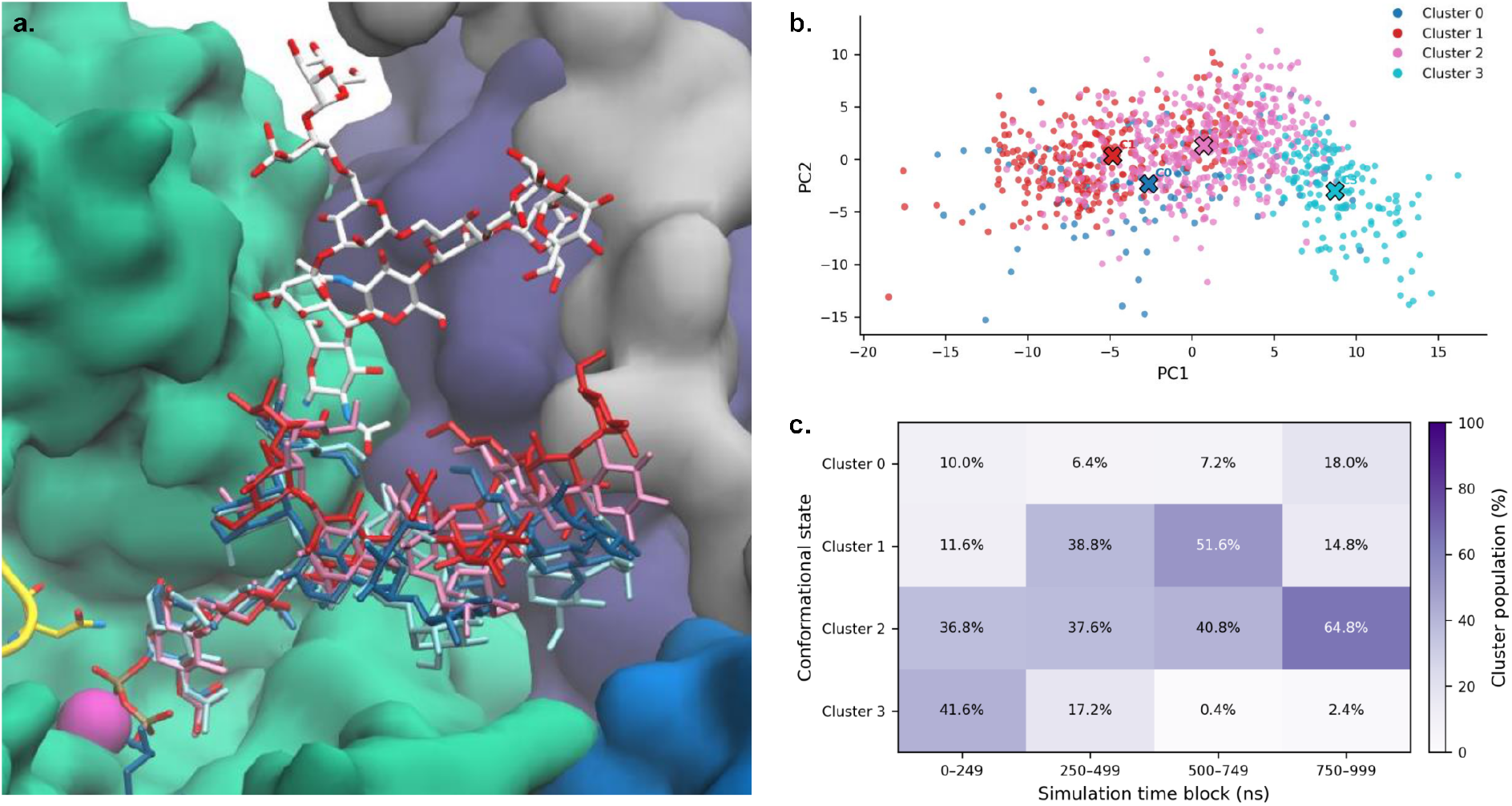
Conformational landscape of the LLO donor glycan during the GaMD simulation. **(a)** Superposition of the four representative LLO donor conformations identified through Gaussian mixture model (GMM) clustering in the OST donor-binding pocket. Protein is rendered as surface with Stt3 in green, Swp1 in grey, Wpb1 in purple. The LLO donor (sticks) is shown with four representative structures (cluster centroids) and coloured accordingly. The N539-linked glycan is shown in sticks, with C atoms in white, N in blue and O in red. Rendering with Visual Molecular Dynamics (VMD) (https://www.ks.uiuc.edu/Research/vmd/). **(b)** Projection of the MD simulation frames onto the first two principal components (PC1 and PC2), coloured according to their GMM cluster assignment. Crosses indicate the centres of the GMM components in PCA space. **(c)** Heatmap showing the relative population of each conformational cluster across four consecutive simulation time blocks (0–249, 250–499, 500–749 and 750–999 ns).

We then analysed how the increased flexibility of the truncated A-branch altered the productive coordination of the reducing end in the catalytic site. OST catalyses transfer of the donor with inversion of the configuration through an SN2-like displacement mechanism (Lairson *et al*. 2008). The acceptor asparagine Nδ2 atom acts as the nucleophile and approaches the anomeric C1 of the GlcNAc from the side opposite the C1–O bond to the dolichyl-pyrophosphate leaving group (Lizak *et al*. 2013; Ramírez and Locher 2023). Transfer requires both close donor–acceptor proximity and appropriate alignment of the nucleophile with the leaving-group axis. We refer to this set of coordinates as ‘reactive alignment’ and quantify it using two parameters, namely the Nδ2–C1 distance (*d_cat*) and the Nδ2–C1–O1 angle (*a_cat*) (**Fig. 5.a**).

**Figure 5.**
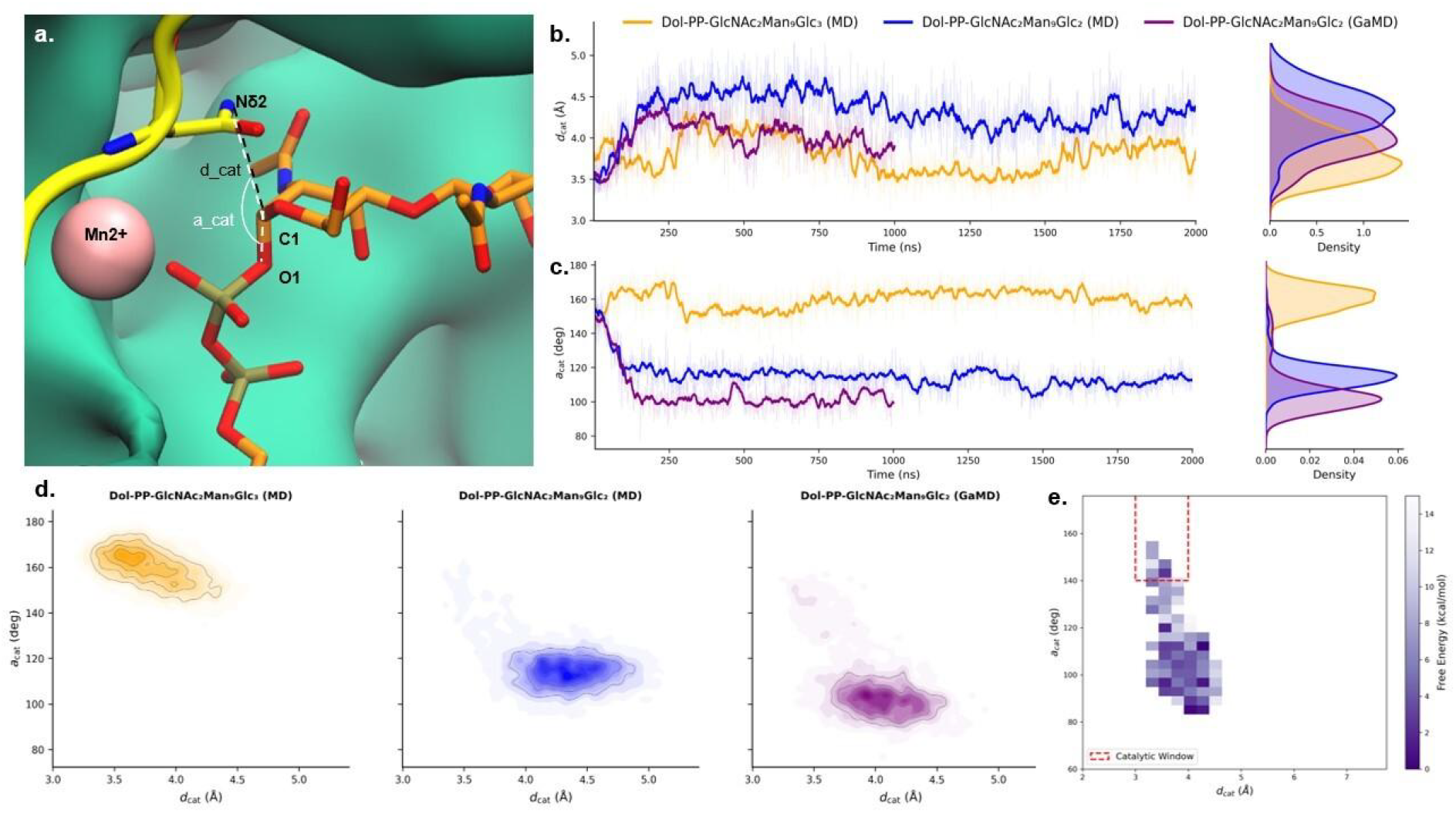
Catalytic geometry of the donor in the OST ternary complex. **(a)** Definition of the donor–acceptor distance, *d_cat*, and nucleophilic attack angle, *a_cat*. **(b and c)** Time series and probability density distributions of *d_cat* **(b)** and *a_cat* **(c)** for the full-length donor in conventional MD (orange) and the truncated donor in conventional MD (blue) and GaMD (purple). **(d)** 2D distributions of *d_cat* and *a_cat* for the three systems. **(e)** Reweighted free-energy surface (FES) of the truncated donor from GaMD, with the catalytically competent region outlined in red. Free energy values are reported relative to the global minimum.

During the deterministic MD sampling, the full-length LLO donor populated a narrow region of the distance–angle space, with the *d_cat* and *a_cat* distributions centred at around 3.8 Å and 160°, respectively, consistent with the near-linear attack geometry expected for an inverting SN2-like reaction (Lairson *et al*. 2008; Schuman, Evans and Fyles 2013) (**Fig. 5.b-d**). By contrast, in both conventional MD and GaMD, the truncated LLO sampled broader, right-shifted distance distributions relative to the optimal alignment, while *a_cat* shifted towards approximately a 100°–120° range, suggesting a loss of optimal reactive alignment (**Fig. 5.b**-**d**). The displacement of the truncated LLO reducing end from the optimal catalytic orientation is clearer when we consider *d_cat* and *a_cat* distributions together in a 2D heat map (**Fig. 5.d**). The full-length LLO populates a compact, high-density region combining short donor–acceptor distances with near-linear attack angles. The position of the truncated LLO is clearly altered, characterised by a non-optimal angular alignment. We can also see that catalytically productive alignments, satisfying both distance and angle requirements(Lairson *et al*. 2008), are rarely sampled by the truncated LLO under both MD sampling conditions.

To assess the relative free-energy accessibility of transfer-compatible configurations in the absence of the terminal glucose, we reconstructed a 2D free energy surface (FES) corresponding to the truncated LLO system by reweighting the GaMD ensemble (**Fig. 5.e**). The results show that the transfer-compatible region, defined here as *d_cat* < 4.0 Å and *a_cat* > 140° and highlighted within a dotted box in red, is only sporadically visited. Instead, the truncated donor was stabilised in low-energy basins characterised by *a_cat* values in a range between 80° and 100°. These basins correspond to the highly populated conformational states identified by PCA clustering (**Fig. 4.a** and **4.b**).

## Conclusions

Most secreted proteins in eukaryotes are N-glycosylated and the efficiency of this process is crucial towards the optimal folding, trafficking, stability and biological function of the large majority of those. OST is the enzyme responsible for initiating N-glycosylation, operating on nascent proteins as they are translocated into the ER lumen. In eukaryotes OST preferentially catalyses the transfer of a complex LLO donor that bears a characteristic glucoside Glc-α(1-2)-Glc-α(1-3)-Glc-α(1-3)-capping the A-branch of the Man9-GlcNAc2-PP-Dol. The terminal Glc-α(1–2)-in this motif, transferred by Alg10 and removed by GI immediately after OST catalysis, has been directly linked to the N-glycosylation efficiency both in yeast(Burda and Aebi 1998; Zacchi and Schulz 2016) and in plants(Farid *et al*. 2011) while defects in ALG10 have been recently linked to congenital disorders in humans affecting the sleep-epilepsy axis(Gill *et al*. 2024). The MD simulations we presented in this work complement this knowledge with a mechanistic framework explaining the preference of eukaryotic OST for a fully glucosylated LLO donor. By combining deterministic sampling and GaMD simulations, we show that the terminal Glc-α(1–2)-anchors the A-branch of the LLO to a distal Wbp1–Ost2 pocket in OST, supporting the stability of the catalytically productive geometry required for glycan transfer. In agreement with previous results and observations(Zacchi and Schulz 2016), we find that the complex structure of the LLO is required to complement the architecture of the OST where the conserved Stt3 N539-linked glycan also contributes to the network of interactions that restricts donor flexibility. These framing contacts extend from the distal Wbp1–Ost2 pocket deep into the catalytic region in Stt3, preserving the coupling between donor–acceptor distance and attack angle required for a transfer-competent state. We show that the removal of the terminal Glc-α(1–2)-disrupts this interaction network and leads to a higher conformational heterogeneity of the truncated LLO. Although a shorter donor is able to form interactions with residues in the pocket and can still transiently approach the acceptor, it does not maintain the optimal catalytic geometry, sampling non-productive conformations.

Taken together, our results show that within the context of the major yeast OST isoform studied here, the terminal Glc-α(1–2)-in the LLO is determinant towards catalytically efficient pre-organisation of the donor, stabilising its position in the binding pocket and preserving the optimal geometry required for catalytic efficiency. Interesting earlier work(Kelleher *et al*. 2003) showed that specific OST isoforms can discriminate between a full LLO donor, ie. Glc3Man9GlcNAc2-PP-Dol, while retaining kinetic transfer competence for assembly intermediates that lack the terminal glucose. Future work on this subject will address how specific changes in the OST architecture complement LLO structure and determine a switch in substrate donor preference.

### Computational Methods

#### Building the system

In our study, we used the cryo-EM structure of the ternary complex involving yeast oligosaccharyltransferase (OST), the lipid-linked oligosaccharide, and a non-acceptor peptide, TAMRA-DAB-NH2 (PDB code: 8AGC) (Ramírez *et al*. 2022) as a starting point. To refine the protein structure, we used SwissModel (Swiss-Model Workspace *et al*.), incorporating the missing residues. The non-acceptor peptide was replaced with the efficiently glycosylated substrate peptide YJR1_N99, previously described and validated in our earlier study. The lipid-linked oligosaccharide (LLO) donor, Glc(α1–2)Glc(α1–3)Glc(α1–3)Man(α1–2)Man(α1–2)Man(α1–3)[Man(α1–2)Man(α1–3)[Man(α 1–2)Man(α1–6)]Man(α1–6)]Man(β1–4)GlcNAc(β1–4)GlcNAc-Dol16, was refined with GlycoShape-3D (Ives *et al*. 2024) (GlyTouCan ID: G91219PM), while the dolichol-16 tail was rebuilt using CHARMM-GUI’s input generator(Jo *et al*. 2008; Lee *et al*. 2016) to ensure CHARMM36m compatibility. The conserved N-glycan linked to N539 of Stt3 was rebuilt as Man8GlcNAc2-as isomer GlyTouCan ID G48369JO with GlycoShape. For both the LLO and the conserved N-glycan, the missing mannose branch was incorporated by pairwise alignment in PyMOL to the cryo-EM-resolved glycans, retaining only the absent branch and adjusting the glycosidic ϕ/ψ angles where necessary according to GlycoShape-3D. The final ternary complex includes the Stt3 (residues 6–682), Ost3 (211–345), Ost2 (22–130), Swp1 (163–283), Wbp1 (263–418) subunits, the peptide substrate (residues –10 to +10 relative to the target asparagine), the dolichyl-diphosphate-linked glycan donor (GlytouCan ID: G91219PM), the conserved N-glycans (GlyTouCan ID: G48369JO) and the catalytic Mn^2+^ metal ion.

#### All-atom molecular dynamics (MD) simulations, deterministic sampling

All-atom MD simulations were performed using version 22 of the AMBER software suite(Wang *et al*. 2004) with the CHARMM36m force field (Best *et al*. 2012; Huang and MacKerell 2013; Vanommeslaeghe and MacKerell 2015). Each system was embedded in a symmetric 130 Å × 130 Å lipid bilayer composed of a 1:1 mixture of POPC lipids (16:0–18:1) using CHARMM-GUI’s input generator(Jo *et al*. 2008; Lee *et al*. 2016) and solvated with TIP3P water(Jorgensen *et al*. 1983). Na^+^ and Cl^−^ ions were added to neutralise the system and reach a final salt concentration of 150 mM. Protein termini were capped (ACE, CT3) and the LLO was modelled using DL16P parameters directly available in CHARMM format. Energy minimization was performed for 5000 steps (2500 steepest descent followed by 2500 conjugate gradient), applying 10 kcal/mol·Å^2^ positional restraints to the protein, LLO, Mn^2+^ ion, peptide substrate, and lipid heads. Long-range dispersion interactions were truncated with a 9 Å cutoff. The systems were gradually heated from 0 K to 310.15 K in three consecutive NVT phases of 125 ps each, with the temperature increased in stages from 0–100 K, 100–200 K, and 200–310.15 K, respectively. Temperature was controlled by Langevin dynamics with collision frequency of 1.0 ps^−1^. The heating phase was followed by a 125 ps NVT equilibration at constant temperature and a 125 ps NPT equilibration at 1 bar using the Berendsen barostat(Berendsen *et al*. 1984) with semi-isotropic pressure coupling. Three further 500 ps NPT equilibration stages were carried out, during which positional restraints were gradually reduced. In the final equilibration step, restraints of 5 kcal/mol·Å^2^ were applied to the protein backbone, the Mn^2+^ ion, and the Cα atom of the targeted Asn, while restraints of 1 kcal/mol·Å^2^ were retained on the peptide substrate and the LLO. Total equilibration time was 2.125 ns. All simulations were run within periodic boundary conditions where long-range electrostatics were treated with Particle Mesh Ewald (PME)(Darden, York and Pedersen 1993) with a 9 Å cutoff. The SHAKE algorithm (Ryckaert, Ciccotti and Berendsen 1977) was used to constrain all bonds to hydrogen atoms, enabling a 2 fs time step. Production MD simulations were performed for 2 μs per system under NPT conditions at 310.15 K and 1 bar. Weak harmonic restraints (5 kcal/mol·Å^2^) were maintained to restrain the glycosylation-site Asn Cα of the peptide in place. The position of the Mn^2+^ ion, of the protein backbone, and of the LLO heavy atoms were also restrained to preserve the integrity of the catalytic architecture within a reduced OST model, while enabling the sampling of the peptide conformation.

#### Hydrogen bonds analysis

Hydrogen bonds were analysed over the 2 μs MD trajectories using the Hydrogen Bonds plugin implemented in VMD (v1.9.4a57). Hydrogen bonds were identified using a donor–acceptor distance cutoff of 4.0 Å and a maximum donor–hydrogen–acceptor angular deviation of 60° from linearity. Hydrogen bond occupancy was calculated as the percentage of analysed trajectory frames in which each interaction was present.

#### Donor LLO dynamics: RMSF analysis

The flexibility of the donor LLO was quantified by calculating the root mean square fluctuation (RMSF) of each glycan residue over the 2 μs molecular dynamics trajectories of both the full and truncated donor systems. Trajectories were processed using MDTraj (v1.10.2) (McGibbon *et al*. 2015) and aligned to the first simulation frame based on the protein Cα atoms to remove overall translational and rotational motions. Atomic RMSF values were calculated with respect to the time-averaged atomic coordinates using NumPy (Harris *et al*. 2020) and grouped by monosaccharide to obtain residue-level distributions. Boxplots were generated using Matplotlib (v3.8.4) (Hunter 2007) in Python.

#### All-atom MD simulations, Gaussian-accelerated (GaMD) sampling

GaMD simulations (Miao, Feher and McCammon 2015; Wang *et al*. 2021) of the OST ternary complex bound to the truncated donor Dol_16_-PP-GlcNAc_2_Man_9_Glc_2_ were performed using the GaMD module in AMBER24 (Case et al., 2024), starting from the same structure used for the deterministic sampling MD simulations.

GaMD was applied in the dual-boost mode, in which independent boost potentials are added to the dihedral and total potential energy terms. The reference energy was set to the lower bound (iE=1), and the upper limit of the standard deviation of the boost potential was fixed at 6 kcal·mol^−1^ for both dihedral (σ0^D^) and total potential (σ0^P^) boosts.

An initial 1 ns conventional MD phase was used to collect potential energy distributions, excluding the first 0.2 ns as pre-processing. These statistics were used by GaMD to determine the optimal acceleration parameters. A 50 ns GaMD equilibration phase was then performed with the boost potential applied. The first 3.2 ns of this phase were excluded from analysis to allow the system to adapt to the modified potential. Finally, a 1 μs GaMD production run was carried out continuously propagating the boost parameters. Potential energy averages and standard deviations were recalculated every 0.5 ns to update the boost parameters during production. All simulations were performed in the NPT ensemble at 310 K and 1 atm, using Langevin dynamics (γ = 1 ps^−1^) for temperature control and isotropic Berendsen barostatting. Hydrogen bonds were constrained with SHAKE (ntc=2, ntf=2), allowing a 2 fs timestep. Long-range electrostatics were evaluated using PME with a 9 Å cutoff for non-bonded interactions. Harmonic restraints (5 kcal·mol^−1^·Å^−2^) were applied to maintain the protein backbone, the catalytic Mn^2+^ ion, and the peptide Asn residue in the active site during the initial phases, matching those used in the conventional sampling MD simulations.

#### Conformational analysis of the truncated donor

GaMD trajectories were analysed to characterise the conformational space of the truncated donor (Dol_16_-PP–GlcNAc_2_Man_9_Glc_2_), with particular focus on the terminal GlcNAc residue and the distal glucose moieties. All trajectory frames were first aligned to the protein backbone to remove global translational and rotational motions. Donor conformations were then described using the Cartesian coordinates of heavy atoms of the selected residues. Coordinates were converted to Å and reshaped into frame-wise feature vectors. The resulting data were standardised (zero mean and unit variance) and subjected to principal component analysis (PCA) using the scikit-learn library (Pedregosa *et al*. 2012). The number of retained principal components was selected to capture at least 95% of the total variance. Clustering was performed in the resulting PCA space using Gaussian mixture models (GMMs) with full covariance matrices (Bishop 2006) as implemented in scikit-learn (Pedregosa *et al*. 2012). Models containing between 1 and 8 components were evaluated, and the optimal number of clusters was selected based on minimisation of the Bayesian Information Criterion (BIC)(Schwarz 1978). Each frame was assigned to the most probable cluster. To evaluate temporal redistribution, the trajectory was divided into four equal time blocks, and cluster populations were computed for each block.

#### Definition of catalytic distance and attack angle

Productive glycan transfer by OST requires nucleophilic attack of the acceptor asparagine side chain on the anomeric carbon of the reducing-end GlcNAc of the lipid-linked oligosaccharide donor. To characterise this geometry, we monitored two collective variables: the donor–acceptor distance (*d_cat*) and the attack angle (*a_cat*). The catalytic distance was defined as the distance between the nucleophilic nitrogen atom (Nδ2) of the acceptor asparagine side chain and the anomeric carbon (C1) of the donor reducing-end GlcNAc. The attack angle was defined by Nδ2–C1–O, where O is the glycosidic oxygen linking the reducing-end GlcNAc to the pyrophosphate–dolichol moiety, and reports on the alignment of the nucleophile relative to the reaction centre and leaving-group direction. Distances and angles were calculated for all frames of the production trajectories using *cpptraj* (AmberTools). Configurations were classified as catalytically competent when both criteria were simultaneously satisfied, with d_cat ≤ 4.0 Å and a_cat ≥ 140°, consistent with near-inline attack geometries expected for glycosyl transfer reactions (Lairson *et al*. 2008). Probability density distributions were estimated by kernel density estimation (KDE), and joint *d_cat-a_cat* distributions were used to assess coupling between donor–acceptor proximity and reactive alignment.

#### Thermodynamic reweighting and free energy surface reconstruction

To recover the unbiased thermodynamic ensemble from the biased GaMD trajectory, reweighting was performed using the PyReweighting toolkit (Miao *et al*. 2014). A two-dimensional potential of mean force (PMF) was calculated along two collective variables defining the catalytic geometry: the donor–acceptor distance (*d_cat*) and the nucleophilic attack angle (*a_cat*).

Reweighting was carried out using the second-order cumulant expansion (CE2) (Miao, Feher and McCammon 2015), which reliably approximates the exponential average of the boost potential (ΔV) under the condition that the standard deviation of the boost distribution remains below 6 kcal mol^−1^. The resulting PMF was represented as a two-dimensional free energy surface (FES) on a grid with bin sizes of 0.25 Å along *d_cat* and 4.0° along *a_cat*. To minimise statistical noise from poorly sampled transient configurations, bins containing fewer than 10 frames over the 1 μs trajectory were excluded. Free energy values were referenced to the global minimum and truncated at an upper limit of 15 kcal mol^−1^.

## Acknowledgements

We gratefully acknowledge the Research Ireland (RI; previously SFI) Centre for Research Training in Foundations of Data Science (www.data-science.ie) for financial support of BT postgraduate training (18/CRT/6049). The University of Southampton is gratefully acknowledged for generous allocation of computational resources on the High-Performance Computing (HPC) cluster Iridis. We would like to thank Prof Benjamin L. Schulz for insightful comments and suggestions on a draft of the manuscript.

## Data Availability

All MD simulations trajectories are available open access (OA) on https://glycoshape.org/downloads and on Zenodo at https://doi.org/10.5281/zenodo.21393289

